# Species and habitat mapping in two dimensions and beyond. Structure-from-Motion Multi-View Stereo photogrammetry for the Conservation Community

**DOI:** 10.1101/2019.12.16.878033

**Authors:** Leon DeBell, James P. Duffy, Trevelyan J. McKinley, Karen Anderson

## Abstract

Structure-from-Motion Multi View Stereo (SfM-MVS) photogrammetry is a technique by which volumetric data can be derived from overlapping image sets, using changes of an objects position between images to determine its height and spatial structure. Whilst SfM-MVS has fast become a powerful tool for scientific research, its potential lies beyond the scientific setting, since it can aid in delivering information about habitat structure, biomass, landscape topography, spatial distribution of species in both two and three dimensions, and aid in mapping change over time – both actual and predicted. All of which are of strong relevance for the conservation community, whether from a practical management perspective or understanding and presenting data in new and novel ways from a policy perspective.

For practitioners outside of academia wanting to use SfM-MVS there are technical barriers to its application. For example, there are many SfM-MVS software options, but knowing which to choose, or how to get the best results from the software can be difficult for the uninitiated. There are also free and open source software options (FOSS) for processing data through a SfM-MVS pipeline that could benefit those in conservation management and policy, especially in instances where there is limited funding (i.e. commonly within grassroots or community-based projects). This paper signposts the way for the conservation community to understand the choices and options for SfM-MVS implementation, its limitations, current best practice guidelines and introduces applicable FOSS options such as OpenDroneMap, MicMac, CloudCompare, QGIS and speciesgeocodeR. It will also highlight why and where this technology has the potential to become an asset for spatial, temporal and volumetric studies of landscape and conservation ecology.

## Introduction

Relatively new technologies, such as drones (1,2), advances in computational power (3,4), improvements in digital cameras (5), along with classic remote sensing platforms such as kite aerial photography (KAP) and balloons (6,7), have all combined to help create a new opportunity in remote sensing research utilising SfM-MVS photogrammetry. It is these convergent developments that now position SfM-MVS as a cost-effective and democratic tool for conservation research.

SfM-MVS approaches use parallax (i.e. minor displacements in similar images) in conjunction with computer vision techniques, in order to derive 3D structures from 2D data (similarly to how brains use parallax from vision to determine the distance and speed of a moving object). Data such as overlapping digital photographs and GPS/GNSS (Global Positioning System/Global Navigation Satellite System) information are stitched together, adding distance (X and Y) and height (Z) values to pixels (points) in the combined data, producing a “point cloud” (Figure 1). From this, SfM-MVS software is able to output two dimensional orthographic images containing detailed geographical location information, alongside 2.5 dimensional reconstructions (2.5D is used as it relates to the limitations of algorithms to produce true 3D reconstructions from aerial images (8)), such as digital elevation, surface and terrain models (DEM, DSM, DTM) (Figure 2). These data products can be used for mapping habitats (9–11), analysing structure, biomass, topography and change (12–15) but also crucially, we suggest, offer the capability to determine the geographic position of species in both two and three dimensions (16–18), thus being of great value for conservation biologists.

**Figure 1:**
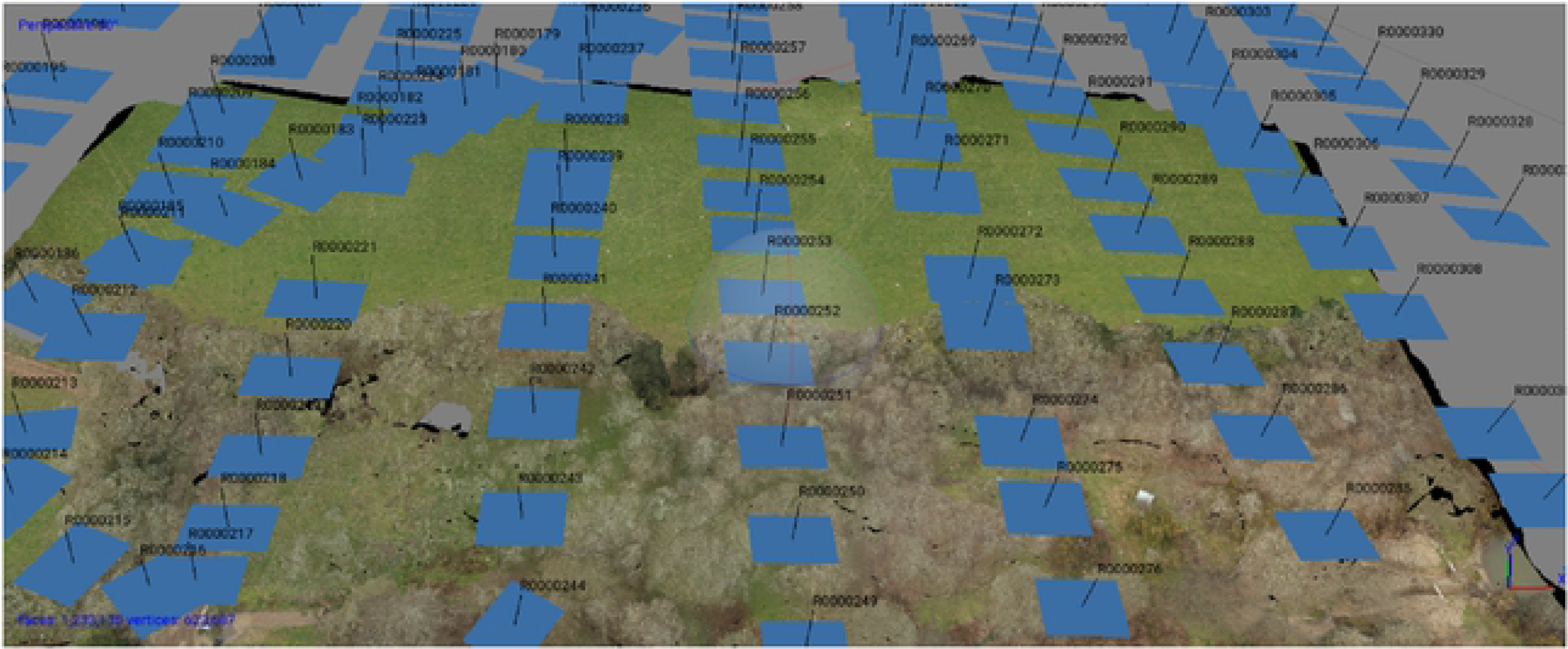
Screenshot from Agisoft Metashape. Blue squares indicate position of the camera in relationship to the ground when images were captured. The height of drone and it’s velocity allow for the same object to be captured from multiple different angles, which in turn allows the SfM-MVS software to calculate the height of objects captured in the scene.

**Figure 2:**
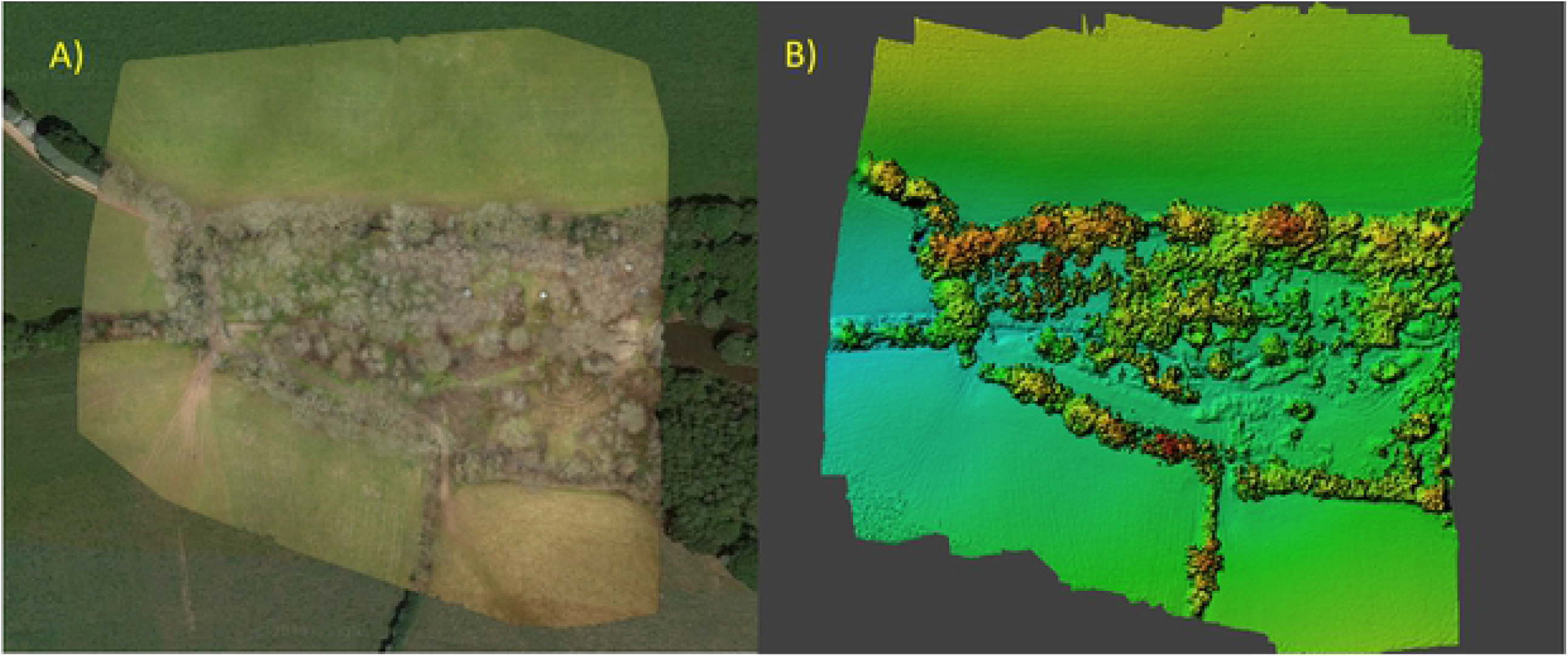
Two typical outputs from SfM-MVS software. A) An orthographic map produced using WebODM. The image was taken from the WebODM interface and shows the produced orthomosaic overlaid on top of the corresponding Google satellite image. B) the DEM produced by Agisoft Metashape. Image taken from within Metashape.

Mapping and modelling the distribution of species is an important aspect of planning and management within conservation; especially with regards to land-use change, and where there is potential for biodiversity loss (19–21). Both Bradbury et al. (21) and Davies and Asner (22) have demonstrated the application of Light Detection and Ranging (LiDAR) for delivering improved spatial understanding of species distributions and predictive modelling. Davies and Answer (22) also demonstrate the importance in understanding habitat in 3D with regards to species populations and communities within ecosystems. Whilst very effective, promoted as a conservation technology by the WWF (23), and even suggested as a tool to archive the Earth in 3D (24), some of the downsides to LiDAR or laser scanning are the cost, weight and lack of portability of instruments, which place this technology out of reach for many with small operating budgets or those working in remote areas (25). In contrast, SfM-MVS techniques have been shown to be a viable alternative and much more cost effective, especially when utilised in conjunction with drone derived data (25–27).

Within the Earth observation sector there has been a recent upsurge in the availability of free and open source analysis tools with which to extract information from those data. One such example is the Google Earth Engine, which allows for easily applied satellite remote sensing techniques with regards to conservation issues (28). Furthermore, there is an expanding range of software and literature that can help in using free and open source software (FOSS) for remote sensing (29–32). Combined with the currently available remote sensing and GIS options, opportunities exist for a completely free and open source SfM-MVS processing pipeline for exploiting two-to three-dimensional data. This paper seeks to address the currently available FOSS SfM-MVS software options, alongside some of the scientifically popular commercial offerings and provide guidance in choice that can be of potential benefit to the conservation community.

## SfM-MVS software ranges and limitations

In recent years, drones have readily captured the media’s attention, but an equal part of their success within a scientific capacity, is due to the software that allows for the processing of the data they collect. Whilst the number of resources discussing the use of drones in conservation is growing (33,34), along with the concepts and processing of SfM-MVS data (33,35) accessible information about implementing SfM-MVS software options can be hard to find or limited. Alongside this, conservation applicable SfM-MVS research tends to be targeted at the geosciences and remote sensing readership, and as such, might not be as accessible or digestible for those in positions of management or decision making within the conservation sphere.

Within the scientific literature, much of the focus is aimed at quantifying outputs from two major commercial software providers; Agisoft and Pix4D. Both of these popular choices only offer discounts to educational institutions and it is likely that their standard prices of $3499 and $4426 (€3990 converted to USD at time of writing) respectively, could easily put them outside of community-based, grassroots or small-scale conservation budgets. Beyond these two options there are now upwards of 40 alternatives available (36), with most of the commercial software costing as much, and in some cases more than Agisoft and Pix4D (Table 1).

**Table 1.** FOSS and sample of popular commercial SfM-MVS options showing operating system compatibility, graphics card requirements, GIS ready outputs, active development and price for perpetual licence. *Prices converted to US$ and correct at time of writing.

| Software | Platform | GPU Required | Native Mapping<br>(orthographic<br>outputs) | Under<br>Development | price for<br>perpetual<br>licence |
| --- | --- | --- | --- | --- | --- |
| Aerial_mapper<br>(37) | Linux | No | Yes | No | \$0 |
| Colmap (38) | Linux,<br>OSX,<br>Windows | Yes (Nvidia only) | No | Yes | \$0 |
| Meshroom (39) | Linux,<br>Windows | Recommended<br>(Nvidia only) | No | Yes | \$0 |
| MicMac (40) | Linux,<br>OSX,<br>Windows | No | Yes | Yes | \$0 |
| OpenDroneMap<br>(41) | Linux,<br>OSX,<br>Windows | No | Yes | Yes | \$0-57 |
| VisualSfM (42) | Linux,<br>OSX,<br>Windows | Recommended<br>(Nvidia only) | No | No | \$0 |
| Regard3D (43) | Linux,<br>OSX,<br>Windows | No | No | Yes | \$0 |
| Agisoft<br>Metashape (44) | Linux,<br>OSX,<br>Windows | Recommended<br>(Nvidia & Radeon) | Yes | Yes | \$3499 |
| Pix4D (45) | Windows | Recommended<br>(Nvidia only) | Yes | Yes | \$4426* |
| RealityCapture<br>(46) | Windows | Yes (Nvidia only) | Yes | Yes | \$16640* |
| DroneMapper<br>(47) | Windows | No | Yes | Yes | \$999 |

Many of the commercial options offer trial periods and we recommend that anyone looking to purchase, utilise the trial period for adequate testing of hardware compatibility and ease of processing. There are many that also offer monthly subscriptions, which can help reduce the burden on initial capital expenditure, but may come with limitations on the number of images available to process, or the amount of data available to transmit or store, especially when the monthly subscription is for a cloud processing service (Table 2). As data collection for SfM-MVS and the subsequent data outputs can quickly produce multiple Gigabytes (GB) of data, cloud or online services can also lead to a requirement for an adequate internet connection, and this should be a consideration when in-situ processing with limited resources is required.

**Table 2:** Comparison/sample of cloud processing options and costs indicating some of the variability of what is available. *Enterprise option with no data restrictions is price on application.

| Software | Monthly cost<br>no contract | Monthly cost<br>with 12-month<br>contract | Data<br>restriction |
| --- | --- | --- | --- |
| Pix4D (45) | \$288 | \$240 | Yes |
| RealityCapture (46) | \$276-\$831 | \$229-693 | Variable |
| DroneDeploy (48) | \$149-POA* | \$99-POA* | Variable |

The available software can be roughly divided into two classes, those that provide survey options (orthographic maps, DEMs and other GIS ready layers), and those that cater more to the recreation of 3D objects via point clouds or meshes. The latter is the result of the growing demand within the gaming and architectural industries and whilst these software have the potential to be utilised, especially where just a point cloud is required, their deployment within the wider conservation community may be more limited, due in part to the extra steps and programmes required to produce data outputs that are likely to be useful in a conservation setting.

Unlike most software used in day to day office work, SfM-MVS software will often leverage all available processing power from a computer, consuming greater levels of energy and generating more heat as a result. If planning to process data in remote locations, it is worth preparing for adequate power supply. Alongside this, there has also been a shift to leveraging graphics cards to process data within certain stages of the SfM-MVS pipeline, and although this can increase processing speeds (Figure 3), it brings with it extra costs associated with necessary hardware. Another potential issue that stems from software that require or benefit from graphics cards, is one whereby not all graphics cards are supported equally, presenting the possibility of owning or purchasing graphics cards that are redundant (Table 1).

**Figure 1.**
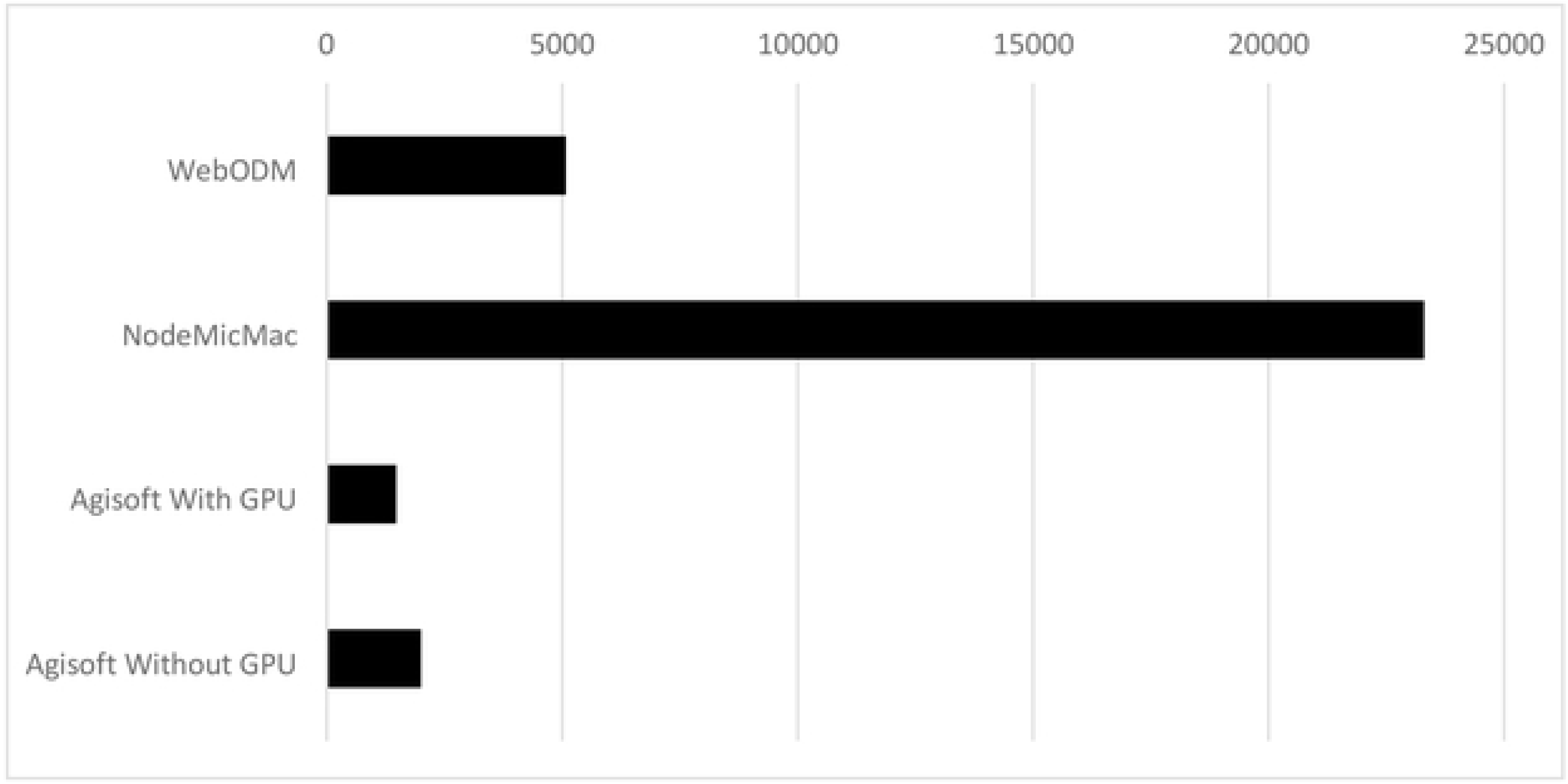
Comparison of processing speeds - data processed for this paper using WebODM, NodeMicMac, Agisoft Metashape without GPU, Metashape with GPU.162JPEG images processed on a desktop running Manjaro Linux OS with AMD Ryzen 2700 CPU, 32GB DDR4 3000Mhz RAM and AMD Vega 56 GPU. It is difficult to replicate exactly the same processing parameters for each software, so defaults were used for WebODM and NodeMicMac, medium settings were used for Agisoft Metashape.

## Open source SfM-MVS

There has been discussion of the impacts that open source hardware technologies such as Arduino and Raspberry Pi could have in helping the conservation industry innovate (49). It is this same open source community that has not only helped to democratise remote sensing (30,50,51), but is also now helping to liberate the production of maps and 3D modelling by building SfM-MVS software. Whilst Aerial Mapper, Colmap, Meshroom, Regard3D and VisualSFM all have potential, when factoring in the ability to easily create orthomosaic maps, DEMs, DTMs or DSMs, alongside point clouds, of the currently available FOSS options (Table 1), this paper will discuss two that are felt to stand out as having extra value within the conservation community; MicMac and OpenDroneMap (ODM).

MicMac is the more mature software of the two and was initially developed in 2003 by the French National Geographic Institute and French national school for geographic sciences (52) whereas ODM began as a concept presentation at the 2014 FOSS4G (Free and Open Source Software for Geospatial) conference in Seoul. Both have drawn attention from the scientific community, but with ODM being the newer option, the focus has primarily been on its development and initial quality testing (3,4,53,54). MicMac’s maturity has allowed for a deeper scrutiny (55,56), has been utilised as the primary SfM-MVS option in geomorphological studies (57) and has been shown to be comparable to both Pix4D and Agisoft Photoscan (Metashape) within a low sward grassland study (58).

Both are also cross platform, meaning that they can be installed on computers running either Linux, Mac OS X or Windows operating systems. ODM also offers “one click” installation options, although these come with a small fee of $57 each but include additional support and a money back guarantee.

There is even now a MicMac “node” that can be used either stand-alone or alongside the ODM web interface, “WebODM”. The MicMac node has been developed by DroneMapper (59), a software and cloud SfM-MVS provider that have built their commercial services upon MicMac. NodeMicMac is still in its infancy but now provides a user two different SfM-MVS processing options from within the one, WebODM interface. Also, as the original version of MicMac can have a steep learning curve (58), this provides an easier, more user-friendly option for deploying MicMac.

With MicMac and ODM being optimised to run without the need for a GPU, they can be installed on almost any computer hardware using a 64bit computer processor within the last 9 years (32) and as such, in situations where conservation project budgets are limited, older, second-hand computer hardware that is available cheaply will suffice.

## Collection of data

Whilst differing in subtle ways, SfM-MVS software all follow a similar processing chain, which has been very well described by Carrivick et al. (35). Creating data products with SfM-MVS depends on the input of data that have been collected with a variety of considerations. Accuracy and precision are not only influenced by the quality and resolution of the sensors used to collect the images, but also by the accuracy and precision of the GNSS receiver providing the spatial information, and where necessary, by the number and position of ground control points (GCPs) utilised (55,60,61). The exception to this is in scenarios whereby actual measurements can be applied to scale a dataset, for example by applying known distances to specific points within a point cloud (62). However, this latter option is only likely useful on smaller scale projects or those that don’t require integration with other geospatial data.

It is important to know what question needs answering beforehand and what scale of error is acceptable. For example, if wanting to produce a fine spatial resolution map, or look at animal species relationship to habitat structure, absolute precision and accuracy might not be as important as it would be with wanting to establish plant below ground biomass derived from above ground biomass (13) or other volume critical measurements (58). Alongside the aforementioned, there are some more useful general principles for data collection which can be applied to any and all SfM-MVS projects;

- Collect data with as high an overlap as is practicable – Nearly every SfM-MVS software will have its own guidelines for the amount of side and forward overlap between images, these range from 60%/60% (63) to 60%/80% (64), though recent studies have shown that higher overlap, upto 90% in both side and forward, is better for accuracy and precision within SfM-MVS derived data (65).
- Collect both nadir and oblique imagery - James and Robson (66) demonstrated that nadir - that is looking straight down - aerial images captured via drone, along with inherent lens distortion of a camera, can induce a systematic error in the form of a doming effect in DEMs. Cunliffe et al. (13) have shown that incorporating oblique imagery can help improve the accuracy of SfM-MVS derived DTMs, whilst Nesbit and Hugenholtz (65) have gone on to quantify that incorporating oblique imagery between 20–35° from nadir, along with higher overlap can improve both precision and accuracy by up to 50%.
- Collect data over a wider area than is required – SfM-MVS data quality deteriorates towards the edges of a reconstruction. Making sure your AOI (area of interest) is central within the data set will ensure there is enough overlap and subsequently be of higher quality (60).
- Cameras with an inbuilt intervalometer simplify data collection – many drones will have either an inbuilt camera mounted on a gimbal, or the ability to trigger a camera via a controller or autopilot, yet if collecting data from a kite or balloon, this will unlikely be possible. So, by choosing a camera that has an intervalometer – the ability to take images at pre-defined time intervals i.e. once every 2 seconds – will simplify data collection. Android apps such as DroneLab Toolkit (67) can also help by providing an intervalometer to cameras in Android phones along with other useful mapping tools.
- Use GCPs – Unless using a camera system or drone equipped with an RTK/PPK (real time or post processing kinematic) GNSS that automatically adds high precision location data into images, the simplest way to adequately warp your model geospatially, is to use GCPs marked using a handheld GNSS (68). For higher accuracy and precision, a DGPS (differential global positioning system) or an RTK/PPK system will be required but can be cost prohibitive. Even with smartphones or cameras that have inbuilt GNSS/GPS and can include this information in images, the use of GCPs can assist in improving the orientation and warping undertaken by the SfM-MVS software and also act as a backup should the GNSS data from a smartphone or camera be of poor quality.
- Time of day – Collecting data around midday, preferably with an overcast/diffuse sky can help improve the overall quality of the data (32). Patchy clouds or bright sunshine, whether early on, or later in the day can all cast shadows which can have an impact when SfM-MVS software stitch the images together.
- Wind – If trying to capture vegetation, especially tall grasses or such that are susceptible to motion from wind, having low or no wind can prove less problematic for SfM-MVS reconstructions. The more stable an object when being captured from multiple positions, the better the possibility for good reconstruction (62,69).
- Setting your camera manually – Mosbrucker et al. (70) give in-depth explanations with regards to relationships between manual settings of cameras, lenses, camera sensors and SfM-MVS, all of which can be important for SfM-MVS outputs. Learn to set a camera manually, starting with obtaining the lowest ISO setting possible, then adjusting aperture, shutter speeds and white balance accordingly (and if possible, manually focused to infinity). This helps to ensure images are taken with consistency in colour, light and subsequently, detail.
- Consideration for camera lens distortion – All consumer cameras have some form of inherent lens distortion (71), which can normally be corrected by SfM-MVS software. However, some cameras, especially sports models have very wide-angle lenses, which create a fish-eye effect. Some SfM-MVS software can automatically correct for this extra distortion, but where not possible FOSS software such as Rawtherapee can be used to correct images before processing (32).
- Collect an extra data set – Forsmoo et al. (58) suggest that, as part of any robust SfM-MVS workflow for ecological purposes, obtaining a replicate data set for comparison can help aid in understanding any error that may arise from the processing through an SfM-MVS pipeline. This could be achieved by collecting twice as many images than is required within the same data capture task.
- Practice and get to understand the software – from our experience, SfM-MVS software will output better or worse data based on slight changes, either in how data is collected, or in how options within the software are utilised. Forums for any given product will usually have answers to problems or can illicit help from those with more experience.

## Processing of Data

Whilst SfM-MVS software typically provide easy to use interfaces for uploading and processing of data, understanding what to expect from the differing SfM-MVS software and understanding what to do with the data once it has been processed can be a challenge. A single dataset can appear differently when processed by differing software (Figure 4), and no software that has been tested is without some form of fault or error (72). As such, in scenarios where precise, centimetre level of error measurements are not required, SfM-MVS software choice can play less of a role (58).

**Figure 4.**
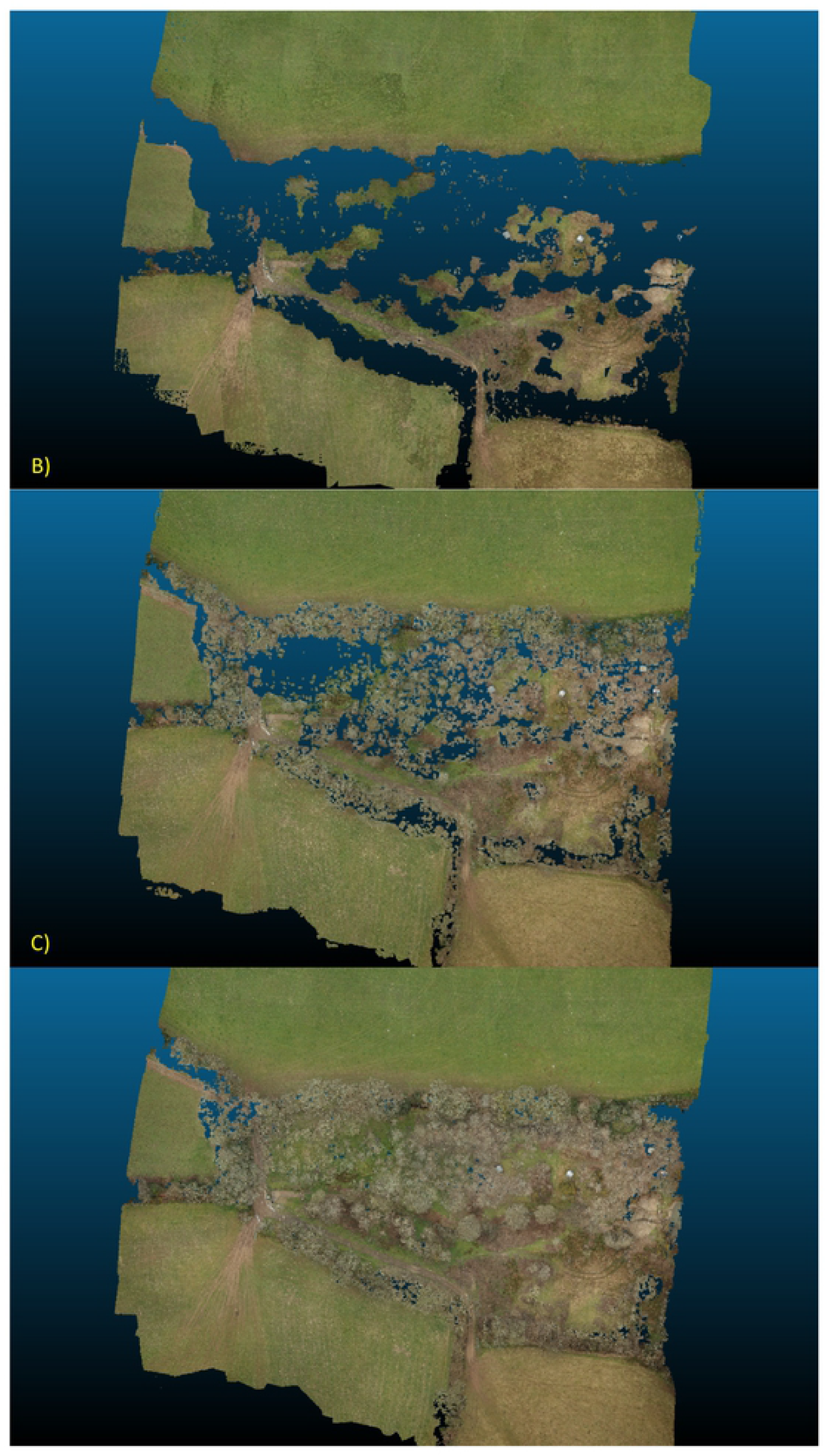
Top down comparison of point clouds produced by WebODM, NodeMicMac and Agisoft Metashape. A) The WebODM pointcloud is devoid of much of the finer detail, with nearly all of the canopy missing from the model. There are potentially several reasons that could cause this but include a result of settings being too aggressive and removing points unnecessarily, or from inadequate overlap in the dataset for ODM to cope with. B) NodeMicMac has retained most of the canopy in the production of its pointcloud, but has excluded much of the ground underneath. C) Agisoft Metashope has produced the most complete pointcloud from the dataset, with both the ground and the canopy being shown and with very few patches of missing data underneath the canopy.

**Figure 5.**
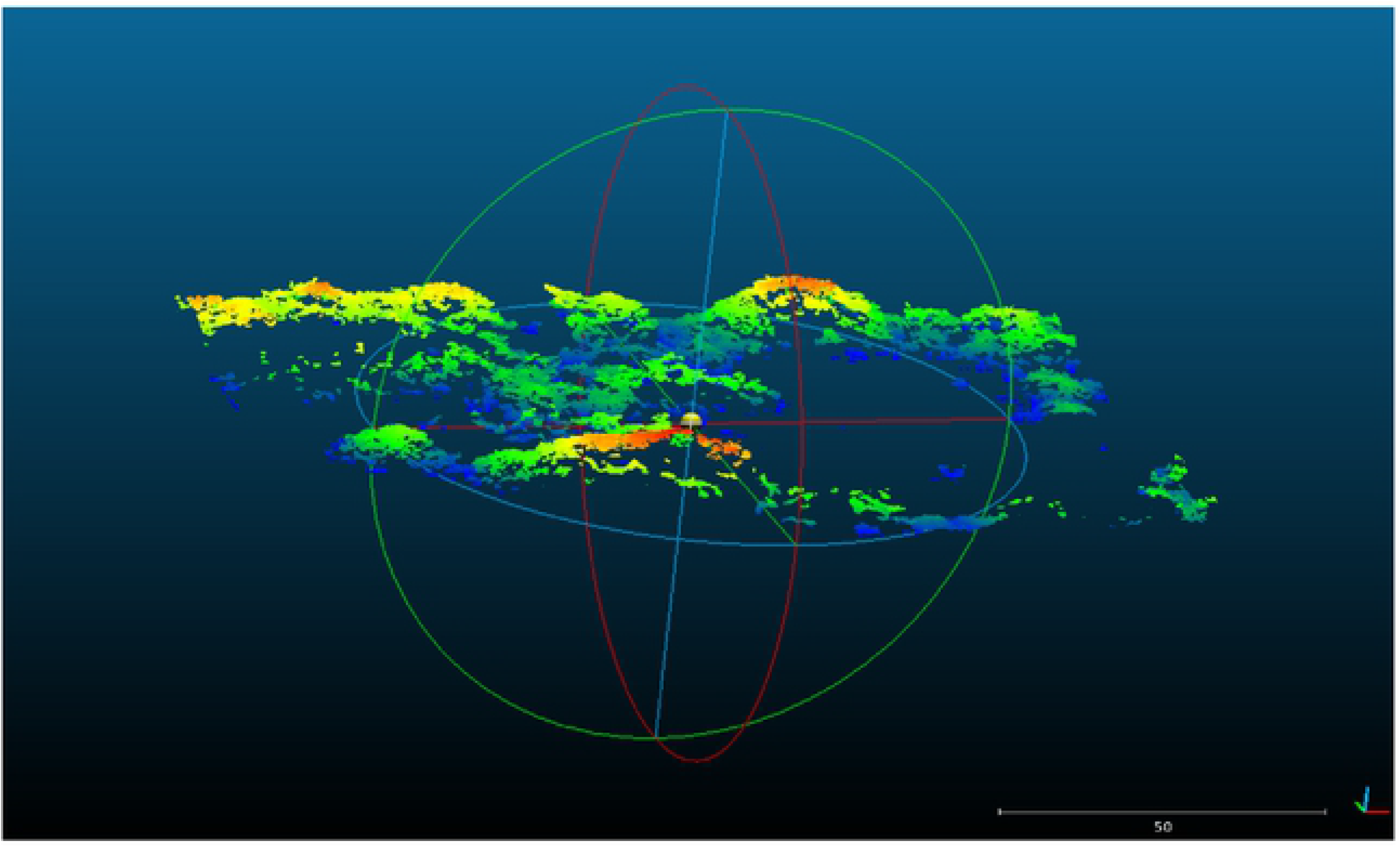
Canopy extracted from the NodeMicMac pointcloud. Using a scalar field on the Z axis within CloudCompare, it is possible to extract the canopy by removing points under a certain height. The canopy can then be examined as a separate entity and labelled if necessary, from with CloudCompare.

Surfaces such as rocks and bare earth can provide an easier medium to recreate, due to their static, exposed nature, whereas vegetation can prove more complicated, with wind possibly affecting reconstruction and canopies possibly being the only source of information, with underlying biomass being masked or blocked.

Many of the software packages have inbuilt features such as distance, area and volumetric measurements, along with point classification (to enable feature examination or extraction based on a type such as a tree, building or road as examples). While these functions may suffice, there are also FOSS geospatial options that can further assist in delivering data products. CloudCompare (73) and QGIS (74) are two such FOSS options that, when combined with modern analysis tools such as the FOSS R-package, speciesgeocodeR (75,76) could prove to be of particular interest within conservation management or policy at any level.

## CloudCompare

This software is dedicated to the reading, manipulation and conversion of 2.5 or 3D data. It offers a multitude of options that include cleaning, filtering, measuring, labelling and cropping of point cloud data along with the ability to perform statistical and computational analysis, both on individual datasets or between multiple datasets.

CloudCompare has proved a useful tool in a number of scientific studies, including those mapping wild leek (77), habitat suitability modelling for insect conservation (78), spatial ecology studies of coral and megabenthic invertebrates (79), and forest canopy ecology (80).

Functions such as its ability to calculate roughness have been used to quantify erosion and deposition of sediment in peatland forests (81) and climate sensitive alpine vegetation (62), whilst change detection via the M3C2 algorithm (82) has been utilised to monitor coral growth (83) and coastal dune morphology (6). The M3C2 function has also proved useful in developing methodologies for quantifying uncertainty in SfM-MVS data (58,61).

## QGIS

QGIS is a software that not only has the ability to process data as a stand-alone GIS desktop application (84), but can also leverage the processing capabilities of other FOSS GIS software such as SAGA (85), GRASS (86), geospatial libraries such as GDAL (87) and Orfeo Toolbox (88), along with programming languages such as R (89) and Python (90).

QGIS has been shown to be useful within conservation research, being used as a main GIS tool in studies such as endemic plant hotspots (91), wildlife management from GPS tracking (92), assessment of conservation area effectiveness with regards to biodiversity and ecosystem services (93) and effects of eutrophication on coral biodiversity (94).

There are a growing number of ‘plugins’ available for QGIS that, when combined with SfM-MVS derived data, could be of benefit to the conservation community too. LecoS (90) adds the ability to automatically analyse the landscape ecology of a raster, the self explanatory ‘Hotspot analysis’ (95), PANDORA for biodiversity ecosystem service assessment (96) and CLUZ (97) for designing conservation and ecological networks. Orfeo Toolbox, although initially a stand-alone programme, can also now be integrated into QGIS via a plugin, enabling a user to perform land cover analysis such as vegetation structure and pattern (98) and assist in conservation planning (99).

A study by Selgrath et al. (100) compared local knowledge with satellite remote sensing for conservation mapping of coral reefs, with QGIS and R serving as the software to process the local knowledge data for map production. It was found that whilst the local environmental knowledge proved useful, where accuracy and precision were considerations, satellite remote sensing still proved the better option, but with a cost approximately 5 times greater.

Utilising SfM-MVS photogrammetry is an opportunity to bridge one such gap by providing both highly detailed maps at a low cost (Figure 6). Freely available satellite images may not be of a high enough resolution, whilst proprietary finer resolution satellite images can be expensive. It can also offer the opportunity for local communities to be involved in the data collection of their local environment.

**Figure 6.**
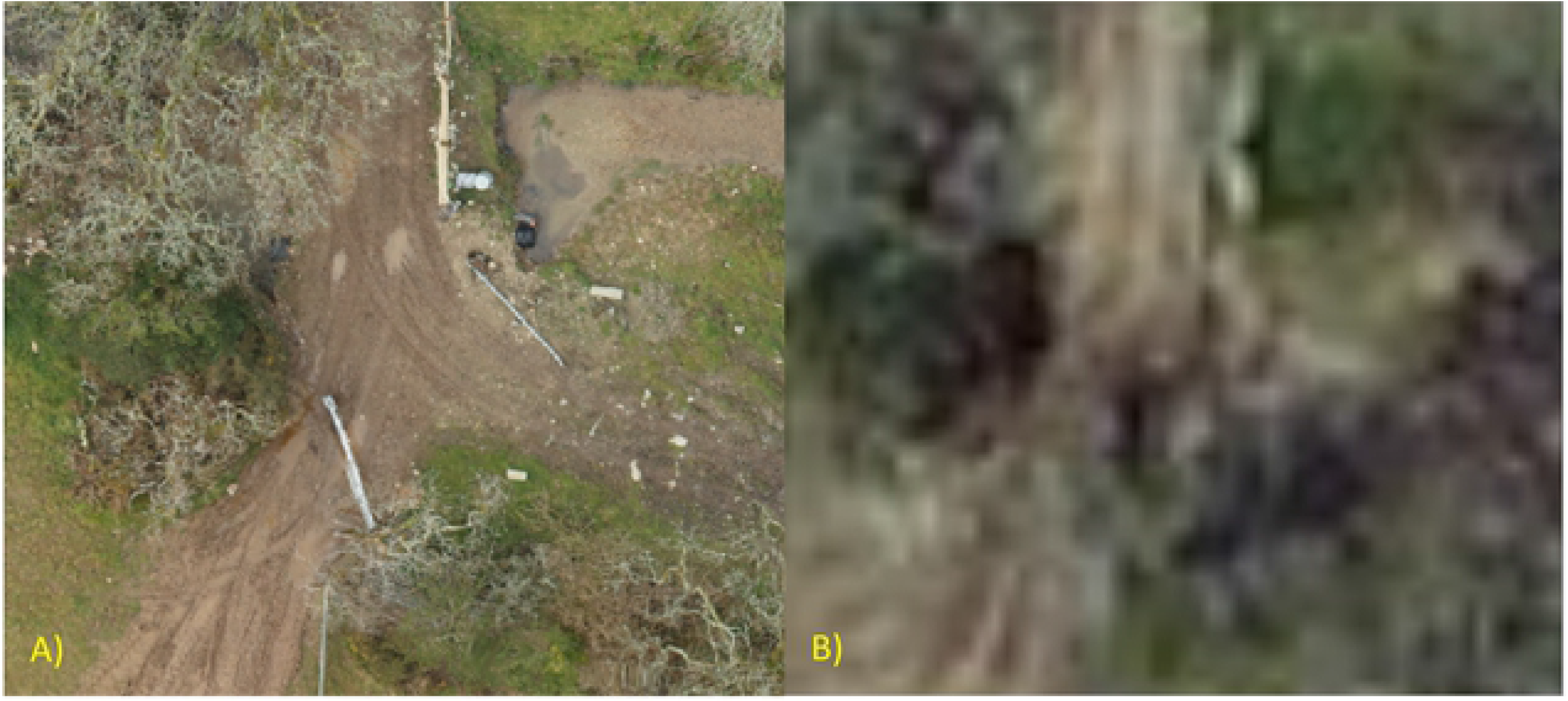
Using QGIS to retrieve the Bing satellite image of the same location, this shows a comparison of resolution between the orthomosaic produced by A) WebODM and B) Bing satellite image. The WebODM orthomosaic image has a resolution of approximately 5cm per pixel, compared to the approximate 50cm per pixel of the Bing satellite image.

## SpeciesgeocodeR

This tool is a relative newcomer to the scientific community and research conducted with it to date has initially focused on the neotropics (101–104). It has been primarily designed for use with big data, dealing with the evolution, biogeography and dispersal of thousands of species across large units of space and time (75,76).

However, with the ability to utilise height/elevation data and by the conclusion presented by Töpel et al. (75) that “the output obtained could be readily used for calculating measures of alpha, beta, and gamma diversity; the identification of neglected areas for conservation; and providing real-time detection of GPS-tagged animals entering and leaving protected areas”, these uses could even be expanded and there exists an opportunity for conservation managers and policy makers to collate and present quantified data in new ways. By combining the high detail outputs of local landscapes from SfM-MVS software, with species data processed by speciesgeocodeR, SfM-MVS can play a crucial role in establishing existing baselines, tracking change and highlighting success and areas for improvement to staff, communities or stakeholders.

## Conclusion

Whether commercial or FOSS, SfM-MVS software offer an opportunity for the conservation community to engage in new and novel remote sensing techniques. They can be utilised to create highly detailed maps and 2.5/3D data at local scales, now include FOSS options that are being successfully deployed for scientific research (3,4,53–58,62,105) and can be incorporated into the existing FOSS remote sensing pipeline. With applications for Android smartphones such as the UAV Toolkit (67) or small, lightweight action cameras now available, combined with balloons, kites, drones, or even collecting data on the ground, there is an opportunity within the conservation sphere to start enabling remote sensing even at a grass-roots or community level, or in any situation where funding may be limited but this type of data desired. In support of the conclusion presented by Berger-Tal and Lahoz-Monfort (49), SfM-MVS as part of a FOSS remote sensing pipeline is one such technology that can help the Conservation industry innovate and dismantle the neo-liberal conservation paradigm. Data of this kind not only opens up opportunities to increase understanding within conservation management, but also has the potential to increase impact when dealing in policy. Further, it presents the opportunity to increase engagement and empowerment within local communities, by providing the ability to collect and view data in-situ for a given project (106,107), whilst providing conservation managers and policy makers the ability to collect and present data in ways that were difficult or overly expensive before. Furthermore, with data visualisation being a powerful medium for communication within ecology (108), the extra options for data visualisation from SfM-MVS could be a valuable tool to better inform stakeholders and engage them in pressing conservation issues and cutting-edge science.

## References

1. Anderson K, Gaston KJ. Lightweight unmanned aerial vehicles will revolutionize spatial ecology. Frontiers in Ecology and the Environment. 2013 Apr 1;11(3):138–46.

2. Ogden LE. Drone Ecology. BioScience. 2013 Sep 1;63(9):776–776.

3. Meza J, Marrugo AG, Ospina G, Guerrero M, Romero LA. A Structure-from-Motion Pipeline for Generating Digital Elevation Models for Surface-Runoff Analysis. J Phys: Conf Ser. 2019 Jun;1247:012039.

4. Myers DJ, Schwelk CM, Wicks R, Bowlick F, Carullo M. Developing A Land Cover Classification Of Salt Marshes Using Uas Time-Series Imagery And An Open Source Workflow. International Archives of the Photogrammetry, Remote Sensing & Spatial Information Sciences. 2018;42(XLII-4/W8):155–62.

5. Crutsinger GM, Short J, Sollenberger R. The future of UAVs in ecology: an insider perspective from the Silicon Valley drone industry. J Unmanned Veh Sys. 2016 Jan 15;4(3):161–8.

6. Duffy J, Shutler J, Witt M, DeBell L, Anderson K. Tracking fine-scale structural changes in coastal dune morphology using kite aerial photography and uncertainty-assessed structure-from-motion photogrammetry. Remote Sensing. 2018;10(9):1494.

7. Feurer D, Planchon O, Maaoui MAE, Ben Slimane A, Boussema MR, Pierrot-Deseilligny M, et al. Using kites for 3-D mapping of gullies at decimetre-resolution over several square kilometres: a case study on the Kamech catchment, Tunisia. Natural Hazards and Earth System Sciences. 2018 Jun 7;18(6):1567–82.

8. Amhar F, Jansa J, Ries C. The Generation Of True Orthophotos Using A 3d Building Model In Conjunction With A Conventional Dtm. 1998;32:7.

9. Fraser RH, Olthof I, Lantz TC, Schmitt C. UAV photogrammetry for mapping vegetation in the low-Arctic. Arctic Science. 2016 Jun 20;2(3):79–102.

10. Palma M, Rivas Casado M, Pantaleo U, Cerrano C. High Resolution Orthomosaics of African Coral Reefs: A Tool for Wide-Scale Benthic Monitoring. Remote Sensing. 2017 Jul;9(7):705.

11. Alexander C, Korstjens AH, Hankinson E, Usher G, Harrison N, Nowak MG, et al. Locating emergent trees in a tropical rainforest using data from an Unmanned Aerial Vehicle (UAV). International Journal of Applied Earth Observation and Geoinformation. 2018 Oct 1;72:86–90.

12. Bareth G, Bolten A, Hollenberg J, Aasen H, Burkart A, Schellberg J. Feasibility study of using non-calibrated UAV-based RGB imagery for grassland monitoring: Case study at the Rengen Long-term Grassland Experiment (RGE), Germany. DGPF Tagungsband 24. 2015;55–62.

13. Cunliffe AM, Brazier RE, Anderson K. Ultra-fine grain landscape-scale quantification of dryland vegetation structure with drone-acquired structure-from-motion photogrammetry. Remote Sensing of Environment. 2016;183:129–143.

14. Mercer JJ, Westbrook CJ. Ultrahigh-resolution mapping of peatland microform using ground-based structure from motion with multiview stereo. Journal of Geophysical Research: Biogeosciences. 2016 Nov 1;121(11):2901–16.

15. Varela MR, Patrício AR, Anderson K, Broderick AC, DeBell L, Hawkes LA, et al. Assessing climate change associated sea-level rise impacts on sea turtle nesting beaches using drones, photogrammetry and a novel GPS system. Global change biology. 2019;25(2):753–762.

16. Wich S, Dellatore D, Houghton M, Ardi R, Koh LP. A preliminary assessment of using conservation drones for Sumatran orang-utan (Pongo abelii) distribution and density. Journal of Unmanned Vehicle Systems Virtual Issue. 2015 Nov 9;01(01):45–52.

17. Cruzan MB, Weinstein BG, Grasty MR, Kohrn BF, Hendrickson EC, Arredondo TM, et al. Small unmanned aerial vehicles (micro-UAVs, drones) in plant ecology. Appl Plant Sci [Internet]. 2016 Sep [cited 2019 Jul 27];4(9). Available from: https://europepmc.org/articles/PMC5033362/

18. McDowall P, Lynch HJ. Ultra-Fine Scale Spatially-Integrated Mapping of Habitat and Occupancy Using Structure-From-Motion. PLOS ONE. 2017 Jan 11;12(1):e0166773.

19. Cowley MJR, Wilson RJ, León-Cortés JL, Gutiérrez D, Bulman CR, Thomas CD. Habitat-based statistical models for predicting the spatial distribution of butterflies and day-flying moths in a fragmented landscape. Journal of Applied Ecology. 2000;37(s1):60–72.

20. Donal PF, Gree RE, Heath MF. Agricultural intensification and the collapse of Europe’s farmland bird populations. Proc Biol Sci. 2001 Jan 7;268(1462):25–9.

21. Bradbury RB, Hill RA, Mason DC, Hinsley SA, Wilson JD, Balzter H, et al. Modelling relationships between birds and vegetation structure using airborne LiDAR data: a review with case studies from agricultural and woodland environments. Ibis. 2005;147(3):443–52.

22. Davies AB, Asner GP. Advances in animal ecology from 3D-LiDAR ecosystem mapping. Trends in Ecology & Evolution. 2014 Dec 1;29(12):681–91.

23. WWF. Conservation Technology [Internet]. WWF. 2019 [cited 2019 Oct 24]. Available from: https://www.wwf.org.uk/project/conservationtechnology

24. Fisher C, Leisz S. The Earth Archive [Internet]. The Earth Archive. 2019 [cited 2019 Dec 11]. Available from: https://www.theeartharchive.com/about

25. Mlambo R, Woodhouse IH, Gerard F, Anderson K. Structure from Motion (SfM) Photogrammetry with Drone Data: A Low Cost Method for Monitoring Greenhouse Gas Emissions from Forests in Developing Countries. Forests. 2017 Mar;8(3):68.

26. Wallace L, Lucieer A, Malenovský Z, Turner D, Vopĕnka P. Assessment of Forest Structure Using Two UAV Techniques: A Comparison of Airborne Laser Scanning and Structure from Motion (SfM) Point Clouds. Forests. 2016 Mar;7(3):62.

27. Glendell M, McShane G, Farrow L, James MR, Quinton J, Anderson K, et al. Testing the utility of structure-from-motion photogrammetry reconstructions using small unmanned aerial vehicles and ground photography to estimate the extent of upland soil erosion. Earth Surface Processes and Landforms. 2017;42(12):1860–1871.

28. Gorelick N, Hancher M, Dixon M, Ilyushchenko S, Thau D, Moore R. Google Earth Engine: Planetary-scale geospatial analysis for everyone. Remote Sensing of Environment. 2017 Dec 1;202:18–27.

29. Pettorelli N, Nagendra H, Williams R, Rocchini D, Fleishman E. A new platform to support research at the interface of remote sensing, ecology and conservation. Remote Sensing in Ecology and Conservation. 2015;1(1):1–3.

30. Wegmann M, Leutner B, Dech S. Remote Sensing and GIS for Ecologists: Using Open Source Software. Exeter: Pelagic Publishing Ltd; 2016. 505 p.

31. OSGeo. Resources Archive [Internet]. OSGeo. 2019 [cited 2019 May 18]. Available from: https://www.osgeo.org/resources/

32. Toffanin P. OpenDroneMap: The Missing Guide. first. MasseranoLabs LLC; 2019.

33. Wich SA, Koh LP. Conservation Drones: Mapping and Monitoring Biodiversity [Internet]. Oxford University Press; 2018 [cited 2019 May 7]. Available from: https://www.oxfordscholarship.com/view/10.1093/oso/9780198787617.001.0001/oso-9780198787617

34. Jiménez López J, Mulero-Pázmány M. Drones for Conservation in Protected Areas: Present and Future. Drones. 2019 Mar;3(1):10.

35. Carrivick J, Smith M, Quincey D. Structure from motion in the geosciences [Internet]. Wiley-Blackwell; 2016. 208 p. Available from: https://www.wiley.com/en-pt/Structure+from+Motion+in+the+Geosciences-p-9781118895849

36. Communicatie FM&. Geomatching | Photogrammetric Imagery Processing Software [Internet]. 2019 [cited 2019 Jul 28]. Available from: https://geo-matching.com/photogrammetric-imagery-processing-software

37. Hinzmann T. Aerial Mapper [Internet]. 2017 [cited 2019 Dec 11]. Available from: https://github.com/ethz-asl/aerial_mapper

38. Schoenberger JL. COLMAP [Internet]. 2019 [cited 2019 Dec 11]. Available from: https://colmap.github.io/license.html

39. Lanthony Y, Castan F. AliceVision Meshroom [Internet]. AliceVision; 2019 [cited 2019 Dec 11]. Available from: https://github.com/alicevision/meshroom

40. MicMac [Internet]. French National Geographic Institute; National Schools of Geographic Sciences; 2003 [cited 2019 Dec 11]. Available from: https://micmac.ensg.eu/index.php/Accueil

41. Benjamin D, Fitzsimmons S, Gargallo P, Toffanin P. OpenDroneMap [Internet]. 2019 [cited 2019 Dec 11]. Available from: https://github.com/OpenDroneMap

42. Wu C. VisualSFM: A Visual Structure from Motion System - Documentation [Internet]. 2011 [cited 2019 Dec 11]. Available from: http://ccwu.me/vsfm/doc.html

43. Hiestand R. Regard3D [Internet]. 2019 [cited 2019 Dec 11]. Available from: http://www.regard3d.org/

44. Agisoft Metashape [Internet]. St Petersburg, Russia: Agisoft LLC; 2006 [cited 2019 Dec 11]. Available from: https://www.agisoft.com/

45. Pix4D [Internet]. Prilly, Switzerland: Pix4D S.A.; 2019 [cited 2019 Dec 11]. Available from: https://www.pix4d.com/

46. RealityCapture [Internet]. Bratislava, Slovakia: Capturing Reality s.r.o.; 2019 [cited 2019 Dec 11]. Available from: https://www.capturingreality.com/

47. DroneMapper. DroneMapper [Internet]. Cedaredge, Colarado: DroneMapper; 2011 [cited 2019 Jul 28]. Available from: https://dronemapper.com/

48. DroneDeploy [Internet]. San Francisco, California: DroneDeploy; 2019 [cited 2019 Dec 11]. Available from: https://www.dronedeploy.com/?utm_source=google&utm_medium=paid%20search&utm_campaign=brand&utm_term=dronedeploy&industry=&role=&stage=&gclid=CjwKCAiAxMLvBRBNEiwAKhr-nIlJMB2lE4XD0Hb4o5Mb-R031E1SszocvupPqG-Z8d8WW6kQwF6PeRoCCb0QAvD_BwE

49. Berger-Tal O, Lahoz-Monfort JJ. Conservation technology: The next generation. Conservation Letters. 2018;11(6):e12458.

50. Jolma A, Ames DP, Horning N, Mitasova H, Neteler M, Racicot A, et al. Free and Open Source Geospatial Tools for Environmental Modelling and Management. In: Jakeman AJ, Voinov AA, Rizzoli AE, Chen SH, editors. Developments in Integrated Environmental Assessment [Internet]. Elsevier; 2008 [cited 2019 May 7]. p. 163–80. (Environmental Modelling, Software and Decision Support; vol. 3). Available from: http://www.sciencedirect.com/science/article/pii/S1574101X08006108

51. Steiniger S, Hay GJ. Free and open source geographic information tools for landscape ecology. Ecological Informatics. 2009 Sep 1;4(4):183–95.

52. Rupnik E, Daakir M, Pierrot Deseilligny M. MicMac – a free, open-source solution for photogrammetry. Open Geospatial Data, Software and Standards. 2017 Jun 5;2(1):14.

53. Park JW, Jeong HH, Kim JS, Choi CU. Development of Open source-based automatic shooting and processing UAV imagery for Orthoimage Using Smart Camera UAV. Int Arch Photogramm Remote Sens Spatial Inf Sci. 2016 Jun 22;XLI–B7:941–4.

54. Stöcker, C., Nex, F., Koeva, M., Gerke, M., Department of Urban and Regional Planning and Geo-Information Management, UT-I-ITC-PLUS, et al. Uav-based cadastral mapping. In: International Archives of the Photogrammetry, Remote Sensing and Spatial Information Sciences [Internet]. International Society for Photogrammetry and Remote Sensing (ISPRS); 2019 [cited 2019 Jul 28]. Available from: https://research.utwente.nl/en/publications/uavbased-cadastral-mapping(c800e413-3b45-40d9-b860-f88a96c62f8f).html

55. Jaud M, Passot S, Allemand P, Le Dantec N, Grandjean P, Delacourt C. Suggestions to Limit Geometric Distortions in the Reconstruction of Linear Coastal Landforms by SfM Photogrammetry with PhotoScan® and MicMac® for UAV Surveys with Restricted GCPs Pattern. Drones. 2019 Mar;3(1):2.

56. Jaud M, Passot S, Le Bivic R, Delacourt C, Grandjean P, Le Dantec N. Assessing the Accuracy of High Resolution Digital Surface Models Computed by PhotoScan® and MicMac® in Sub-Optimal Survey Conditions. Remote Sensing. 2016 Jun;8(6):465.

57. Girod LMR, Nuth C, Kääb A, Etzelmüller B, Köhler JC. Terrain changes from images acquired on opportunistic flights by SfM photogrammetry. The Cryosphere. 2017;11(2):827–40.

58. Forsmoo J, Anderson K, Macleod CJA, Wilkinson ME, DeBell L, Brazier RE. Structure from motion photogrammetry in ecology: Does the choice of software matter? Ecology and Evolution. 2019;0(0):1–16.

59. JP. NodeMICMAC - a new WebODM Node [Internet]. DroneMapper. 2019 [cited 2019 Jul 28]. Available from: https://dronemapper.com/nodemicmac-a-new-webodm-node/

60. Akturk E, Altunel AO. Accuracy assessment of a low-cost UAV derived digital elevation model (DEM) in a highly broken and vegetated terrain. Measurement. 2019 Mar 1;136:382–6.

61. James MR, Robson S, Smith MW. 3-D uncertainty-based topographic change detection with structure-from-motion photogrammetry: precision maps for ground control and directly georeferenced surveys. Earth Surface Processes and Landforms. 2017;42(12):1769–1788.

62. Niederheiser R, Rutzinger M, Lamprecht A, Steinbauer K, Winkler M, Pauli H. Mapping Alpine Vegetation Location Properties By Dense Matching. Int Arch Photogramm Remote Sens Spatial Inf Sci. 2016 Jun 16;XLI–B5:881–6.

63. DroneDeploy. Mapping varied elevations, tall buildings or trees [Internet]. 2019 [cited 2019 Jul 29]. Available from: https://support.dronedeploy.com/docs/mapping-varied-elevations-tall-buildings-or-trees

64. Agisoft. Useful Tips on Image Capture - How to Get an Image Dataset that Meets Metashape Requirements? [Internet]. Agisoft; 2019 [cited 2019 Jul 29]. Available from: https://www.agisoft.com/pdf/tips_and_tricks/Image%20Capture%20Tips%20-%20Equipment%20and%20Shooting%20Scenarios.pdf

65. Nesbit PR, Hugenholtz CH. Enhancing UAV–SfM 3D Model Accuracy in High-Relief Landscapes by Incorporating Oblique Images. Remote Sensing. 2019 Jan;11(3):239.

66. James MR, Robson S. Mitigating systematic error in topographic models derived from UAV and ground-based image networks. Earth Surface Processes and Landforms. 2014;39(10):1413–20.

67. Anderson K, Griffiths D, DeBell L, Hancock S, Duffy JP, Shutler JD, et al. A grassroots remote sensing toolkit using live coding, smartphones, kites and lightweight drones. PloS one. 2016;11(5):e0151564.

68. Cao C, Lee X, Xu J. Positional and Dimensional Accuracy Assessment of Drone Images Geo-referenced with Three Different GPSs. In: AGU Fall Meeting Abstracts. 2017. p. A33I–2499.

69. Dandois JP, Olano M, Ellis EC. Optimal Altitude, Overlap, and Weather Conditions for Computer Vision UAV Estimates of Forest Structure. Remote Sensing. 2015 Oct;7(10):13895–920.

70. Mosbrucker AR, Major JJ, Spicer KR, Pitlick J. Camera system considerations for geomorphic applications of SfM photogrammetry. Earth Surface Processes and Landforms. 2017;42(6):969–86.

71. Ingwer P, Ingwer P, Gassen F, Stefan P, Duhn M, Sch M, et al. Practical usefulness of structure from motion (SfM) point clouds obtained from different consumer cameras. Proceedings of SPIE - Mobile Devices and Multimedia: Enabling Technologies, Algorithms, and Applications. 2015;9411(March):1–11.

72. Remondino F, Nocerino E, Toschi I, Menna F. A Critical Review Of Automated Photogrammetric Processing Of Large Datasets. 2017;9.

73. Girardeau-Montaut D. CloudCompare - Open Source project [Internet]. 2019 [cited 2019 Dec 11]. Available from: https://www.danielgm.net/cc/

74. Sherman G. QGIS [Internet]. 2002 [cited 2019 Dec 11]. Available from: https://qgis.org/en/site/index.html

75. Töpel M, Zizka A, Calió MF, Scharn R, Silvestro D, Antonelli A. SpeciesGeoCoder: Fast Categorization of Species Occurrences for Analyses of Biodiversity, Biogeography, Ecology, and Evolution. Syst Biol. 2017 Mar 1;66(2):145–51.

76. Zizka A, Antonelli A. speciesgeocodeR: An R package for linking species occurrences, user-defined regions and phylogenetic trees for biogeography, ecology and evolution. bioRxiv. 2015 Nov 24;032755.

77. Leduc M-B, Knudby AJ. Mapping Wild Leek through the Forest Canopy Using a UAV. Remote Sensing. 2018 Jan;10(1):70.

78. Stephenson N, Perroy R, Eiben J, Klasner F. High resolution habitat suitability modelling for an endemic restricted-range Hawaiian insect (Nysius wekiuicola, Hemiptera: Lygaeidae). J Insect Conserv. 2017 Feb 1;21(1):87–96.

79. Robert K, Huvenne VAI, Georgiopoulou A, Jones DOB, Marsh L, Carter GDO, et al. New approaches to high-resolution mapping of marine vertical structures. Sci Rep. 2017 Aug 21;7(1):1–14.

80. Hess C, Härdtle W, Kunz M, Fichtner A, Oheimb G von. A high-resolution approach for the spatiotemporal analysis of forest canopy space using terrestrial laser scanning data. Ecology and Evolution. 2018;8(13):6800–11.

81. Stenberg L, Tuukkanen T, Finér L, Marttila H, Piirainen S, Kløve B, et al. Evaluation of erosion and surface roughness in peatland forest ditches using pin meter measurements and terrestrial laser scanning. Earth Surface Processes and Landforms. 2016;41(10):1299–311.

82. Lague D, Brodu N, Leroux J. Accurate 3D comparison of complex topography with terrestrial laser scanner: Application to the Rangitikei canyon (N-Z). ISPRS Journal of Photogrammetry and Remote Sensing. 2013 Aug 1;82:10–26.

83. Neyer F, Nocerino E, Gruen A. Monitoring Coral Growth - The Dichotomy Between Underwater Photogrammetry And Geodetic Control Network. Int Arch Photogramm Remote Sens Spatial Inf Sci. 2018 May 30;XLII–2:759–66.

84. Brovelli MA, Minghini M, Moreno-Sanchez R, Oliveira R. Free and open source software for geospatial applications (FOSS4G) to support Future Earth. International Journal of Digital Earth. 2017 Apr 3;10(4):386–404.

85. Passy P, Théry S. The Use of SAGA GIS Modules in QGIS. In: QGIS and Generic Tools [Internet]. John Wiley & Sons, Ltd; 2018 [cited 2019 Oct 8]. p. 107–49. Available from: https://onlinelibrary.wiley.com/doi/abs/10.1002/9781119457091.ch4

86. Lacaze B, Dudek J, Picard J. GRASS GIS Software with QGIS. In: QGIS and Generic Tools [Internet]. John Wiley & Sons, Ltd; 2018 [cited 2019 Oct 8]. p. 67–106. Available from: https://onlinelibrary.wiley.com/doi/abs/10.1002/9781119457091.ch3

87. Ose K. Introduction to GDAL Tools in QGIS. In: QGIS and Generic Tools [Internet]. John Wiley & Sons, Ltd; 2018 [cited 2019 Oct 8]. p. 19–65. Available from: https://onlinelibrary.wiley.com/doi/abs/10.1002/9781119457091.ch2

88. Cresson R, Grizonnet M, Michel J. Orfeo ToolBox Applications. In: QGIS and Generic Tools [Internet]. John Wiley & Sons, Ltd; 2018 [cited 2019 Oct 8]. p. 151–242. Available from: https://onlinelibrary.wiley.com/doi/abs/10.1002/9781119457091.ch5

89. Muenchow J, Schratz P, Brenning A. RQGIS: Integrating R with QGIS for Statistical Geocomputing. The R Journal. 2017;9(2):409.

90. Jung M. LecoS — A python plugin for automated landscape ecology analysis. Ecological Informatics. 2016 Jan 1;31:18–21.

91. Cañadas EM, Fenu G, Peñas J, Lorite J, Mattana E, Bacchetta G. Hotspots within hotspots: Endemic plant richness, environmental drivers, and implications for conservation. Biological Conservation. 2014 Feb 1;170:282–91.

92. Cagnacci F, Urbano F. Managing wildlife: A spatial information system for GPS collars data. Environmental Modelling & Software. 2008 Jul 1;23(7):957–9.

93. Márquez JRG, Krueger T, Páez CA, Ruiz-Agudelo CA, Bejarano P, Muto T, et al. Effectiveness of conservation areas for protecting biodiversity and ecosystem services: a multi-criteria approach. International Journal of Biodiversity Science, Ecosystem Services & Management. 2017 Jan 1;13(1):1–13.

94. Duprey NN, Yasuhara M, Baker DM. Reefs of tomorrow: eutrophication reduces coral biodiversity in an urbanized seascape. Global Change Biology. 2016;22(11):3550–65.

95. Oxoli D, Zurbarán MA, Shaji S, Muthusamy AK. Hotspot analysis: a first prototype Python plugin enabling exploratory spatial data analysis into QGIS [Internet]. PeerJ Inc.; 2016 Sep [cited 2019 Oct 9]. Report No.: e2204v4. Available from: https://peerj.com/preprints/2204

96. Pelorosso R, Gobattoni F, Geri F, Leone A. PANDORA 3.0 plugin: A new biodiversity ecosystem service assessment tool for urban green infrastructure connectivity planning. Ecosystem Services. 2017 Aug 1;26:476–82.

97. Smith R. The CLUZ plugin for QGIS: designing conservation area systems and other ecological networks. Research Ideas and Outcomes. 2019 Jan 31;5:e33510.

98. Adjonou K, Kémavo A, Fontodji JK, Tchani W, Sodjinou F, Sebastià MT, et al. Vegetation dynamics patterns, biodiversity conservation and structure of forest ecosystems in the wildlife reserve of Togodo in Togo, West Africa. 2017 Aug [cited 2019 Oct 9]; Available from: https://repositori.udl.cat/handle/10459.1/60376

99. Alleaume S, Dusseux P, Thierion V, Commagnac L, Laventure S, Lang M, et al. A generic remote sensing approach to derive operational essential biodiversity variables (EBVs) for conservation planning. Methods in Ecology and Evolution. 2018;9(8):1822–36.

100. Selgrath JC, Roelfsema C, Gergel SE, Vincent ACJ. Mapping for coral reef conservation: comparing the value of participatory and remote sensing approaches. Ecosphere. 2016;7(5):e01325.

101. Gomes VHF, Vieira ICG, Salomão RP, Steege H ter. Amazonian tree species threatened by deforestation and climate change. Nat Clim Chang. 2019 Jul;9(7):547–53.

102. Maldonado C, Molina CI, Zizka A, Persson C, Taylor CM, Albán J, et al. Estimating species diversity and distribution in the era of Big Data: to what extent can we trust public databases? Global Ecology and Biogeography. 2015;24(8):973–84.

103. Zizka A. Big data suggest migration and bioregion connectivity as crucial for the evolution of Neotropical biodiversity. Frontiers of Biogeography [Internet]. 2019 [cited 2019 Oct 9];11(2). Available from: https://escholarship.org/uc/item/25h0w1cx

104. Zizka A, Steege H ter, Pessoa M do CR, Antonelli A. Finding needles in the haystack: where to look for rare species in the American tropics. Ecography. 2018 Feb 1;41(2):321–30.

105. Forsmoo J, Anderson K, Macleod CJA, Wilkinson ME, Brazier R. Drone-based structure-from-motion photogrammetry captures grassland sward height variability. Journal of Applied Ecology. 2018;55(6):2587–99.

106. Brown G, Strickland-Munro J, Kobryn H, Moore SA. Stakeholder analysis for marine conservation planning using public participation GIS. Applied Geography. 2016 Feb 1;67:77–93.

107. Strager MP, Rosenberger RS. Incorporating stakeholder preferences for land conservation: Weights and measures in spatial MCA. Ecological Economics. 2006 Jun 1;57(4):627–39.

108. McInerny GJ, Chen M, Freeman R, Gavaghan D, Meyer M, Rowland F, et al. Information visualisation for science and policy: engaging users and avoiding bias. Trends in Ecology & Evolution. 2014 Mar 1;29(3):148–57.

